# A Software for Identification and Characterization of Theta Rhythms in the Hippocampus

**DOI:** 10.1101/2025.03.25.645280

**Authors:** Nooshin Bahador, Spandan Sengupta, Josh Saha, Milad Lankarany, Liang Zhang, Frances K Skinner

## Abstract

Characterizing theta rhythms in the hippocampus provides a window into understanding memory processing. An inquiry that arises when an animal sustains a pathological state is how theta rhythms are affected. In pathological states like epilepsy or Alzheimer’s, these rhythms change in specific ways. Statistically robust changes in these rhythms could serve as potential biomarkers, indicating the severity of the animal’s condition and the effectiveness of a drug. However, this understanding depends on how the data is analyzed. There are currently no standard criteria for recognizing theta dominance in experimental recordings. To address this, we have developed novel MATLAB-based software with an easy-to-use graphical user interface which enables identifying and analyzing theta rhythms in a standard way. We discuss the software’s functionality and its underlying algorithms. The algorithms were developed using previously acquired EEG/LFP data recorded from the hippocampus of a mouse kindling model of epilepsy. Two primary analyses were conducted to test the software’s functionality: first, comparing theta rhythms during the baseline period versus during spontaneous recurrent seizures; second, analyzing the timing of theta rhythms relative to the seizure event. Our illustrative results indicate that our developed software can robustly identify theta events with statistically significant feature differences. Further, the examination presented here with two mice shows that while theta events can occur just before seizures, it takes tens of minutes post-seizure before theta rhythms occur again. Our software thus provides the user with the ability to robustly identify and characterize theta rhythms and their feature changes.

**Significance Statement:** Theta rhythms in the hippocampus are fundamental for spatial navigation and memory formation. Their observed changes during several pathological states such as epilepsy and Alzheimer’s make them highly interesting to be able to serve as biomarkers. However, their variability (in terms of the time of occurrence, duration, and frequency range) makes them challenging to quantify. The absence of available tools for the automatic extraction of these rhythms from extended datasets significantly hampers processing efficiency and reduces accuracy and consistency. Clinicians and researchers often manually inspect their data to identify these rhythms, a process that is not only time-consuming but also inherently subjective. We have thus developed a MATLAB software for precise, automatic analysis of theta rhythms in EEG/LFP recordings.

## 1. Introduction

Theta rhythms (∼4-12 Hz) in the hippocampus are fundamental and were recorded over 80 years ago (Buzsáki, 2002; Colgin, 2013). They are observed during locomotion (running or walking) or exploratory behaviors (sniffing to investigate new environments) and play key roles in encoding and memory representations.

Memory processes can be considered as a sequence of neuronal patterns activating over time. Each pattern comprises neuronal assemblies, collectively forming a cohesive representation of the external world at any moment. This representation includes various aspects of experience like location and sensory inputs, each linked to a unique pattern of brain activity. For instance, place cells, specific pyramidal neurons in the hippocampus, show spatial specificity. These neurons activate when the animal is in a specific location, creating a neural code for physical position. During a behavioural event, the integration of multi-modal information from various sensory inputs into a unified representation is essential, and theta rhythms help link the separate components of an experience. This sequential organization is important because experiences are inherently temporal. Events are encoded as a sequence of representations, preserving their temporal order. Further, theta rhythms are often coupled with faster gamma cycles which may refer to a particular representation. In essence, theta rhythms are essential for spatial navigation and memory formation, helping the brain construct and retrieve memory traces (Colgin, 2013, 2016; Lisman et al, 2017).

Cognitive changes are reflected in the brain’s expression of theta activities. In pathological states like epilepsy, traumatic brain injury or Alzheimer’s, theta rhythms and their coupling to gamma rhythms are known to change in specific ways (e.g., Chauvière et al., 2009; Fedor et al., 2010; Goutagny et al., 2013; Milikovsky et al., 2017). Clear, statistically robust changes in these rhythms could serve as potential biomarkers, indicating the severity of the animal’s condition and the effectiveness of a drug. However, the community does not currently employ standard criteria for defining or recognizing theta dominance in EEG/local field potential (LFP) recordings. Further, a significant challenge lies in the large variability that occurs in recordings during pathological states, making it important to characterize theta rhythm features beyond just frequency. Thus, considering how these rhythms change cannot be well-quantified or compared in a consistent fashion as it depends on how the data is analyzed. Typically, theta rhythms extracted from experimental datasets are dependent on the experimenter’s intuition and observations, and are thus subjective by nature.

To address this, we here present a MATLAB software code with a convenient graphical user interface (GUI) that can identify dominant theta rhythms in EEG/LFP recordings and characterize their features. In this way, one can thoroughly analyze and quantify theta features in a given dataset and statistically compare it with other datasets. We use the pathology of seizures, as brought about in a kindling mouse model to develop and illustrate our software, but the code can be used to extract and analyze dominant theta rhythms in any EEG/LFP recordings.

## 2. Materials and Methods

### 2.1. Experimental Data and Theta Activity Identification Criteria

To consider theta rhythm features and changes associated with pathological states, we used previously acquired EEG/LFP data from a mouse seizure model and its associated controls. The mouse model is a kindling-based one that has been widely used to study epilepsy. In particular, a modified stimulation protocol was developed for an extended kindling approach such that spontaneous recurrent seizures (SRS) could be obtained (Bin et al., 2017). C57 male black mice were used and each animal had two implanted pairs of electrodes: one pair for stimulation and recording, and another pair for recording from an unstimulated structure. The data used as illustration in this paper is from unstimulated middle hippocampal CA3. We had aimed to target the middle hippocampal CA3 for kindling stimulation (Liu et al., 2021). The stereotaxic coordinates of the CA3 electrode implantation were at bregma −2.5 mm, lateral 3.0 mm, and depth 3.0 mm (Franklin and Paxinos, 1997). The tracks of the implanted CA3 electrodes were histologically assessed, and putative electrode tips in the CA3a or CA3b subregion were observed in several mice. Baseline recordings were taken before starting the kindling process so that a comparison of normal and pathological states in the same animal would be possible.

Spatial memory deficits were shown to be present in these kindled animals (Liu et al., 2021; Song et al., 2018, 2024), and theta rhythms were present in both normal and pathological states. However, identifying theta activity was challenging in terms of analyzing many hours of data manually, thus finding time instances when theta rhythms are dominant is even more complicated. This latter challenge was due to a switching between ‘delta’ and ‘theta’ states that often occurred so that finding recordings of sufficient length where theta rhythms were dominant was difficult. Moreover, instances where theta rhythms were dominant were very sparse. Most of the time, theta rhythms were contaminated by either high-power lower or higher frequencies, which prevented them from being considered as dominant. Indeed, the expression of delta dominant rhythms during awake states has been well-characterized (Schultheiss et al., 2020). Considering all of this, we developed an open access code with an appealing user interface to automatically extract dominant theta rhythms from long recordings.

Since 4 Hz frequency marks the lower boundary of the theta band and the upper limit of the delta band, and mice exhibited minimal theta activity at 4 Hz, we chose a threshold of 5 Hz for the lower end of the theta range. For mice, the theta band typically starts from 5-6 Hz, peaking around 8-10 Hz, and extending up to 12-15 Hz. We thus considered that we would determine theta dominant regions in the recordings by identifying a ratio of theta power to delta power that consistently exceeded a given threshold for a given duration. A higher frequency band would also be included for a relative ratio threshold of an upper bound. Taking all of these factors into account, the criteria for selecting dominant theta activity in this study included three frequency bands – a low-frequency (delta) band of 1-4 Hz, a mid-frequency (theta) band of 5-15 Hz, and a higher frequency band of 16-19 Hz. The theta-delta ratio and theta-higher ratio, each greater than 1.5, and a minimum duration of 5 seconds marked what we refer to as a *theta-dominant segment*.

### 2.2. Software Description

The developed software, ***Neuro Dominance Tracker***, is freely available via a public OSF site [https://osf.io/rf3mn/], and has been tested on different MATLAB versions (2023, 2024) and operating systems (windows, linux, apple). Step-by-step instructions are provided in a manual [https://docs.google.com/document/d/10qLwA9YZCf_VvvBvtcigR9_40KGqYdSXEib50vZ26j8/edit?tab=t.0]. **Figure 1** outlines the steps of the software that are described in detail below.

**Figure 1:**
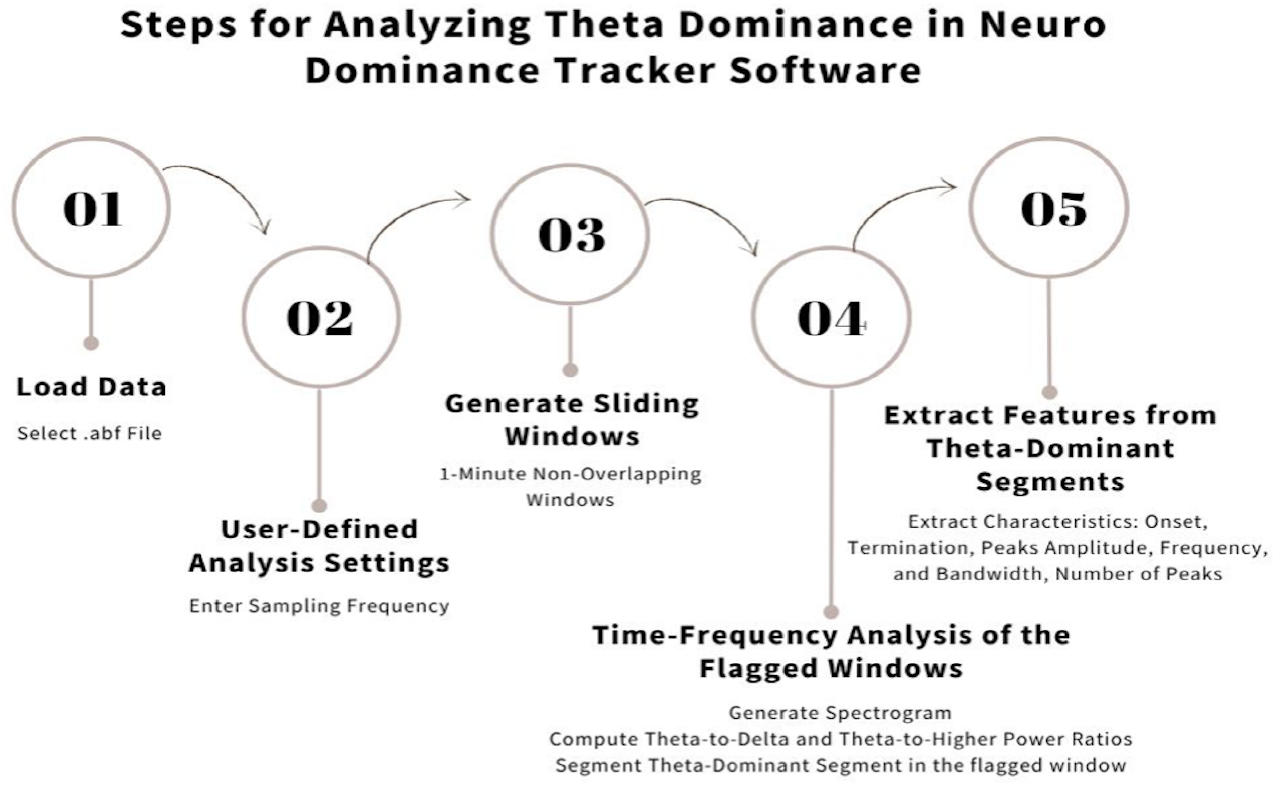
Neuro Dominance Tracker Overview. Steps 01-05 represent the 5 steps that form part of the software to identify and extract theta-dominant segments in EEG/LFP recordings. Overviews of spectrogram generation (section 2.2.4) and identification of theta-dominant segments (section 2.2.5) are provided in *Figure 1-1* in Extended Data.

#### 2.2.1. File Support

By default, the software is configured to load ABF, FIF, EDF, and BDF formats. The user can select the file through a file dialog, and the software utilizes the abfload function as well as other Fieldtrip functions (Oostenveld, 2011) to import the data. data. All needed files are available from the public OSF site. However, the link to access Fieldtrip can be downloaded from https://www.fieldtriptoolbox.org/.

Users can also modify the source code easily to support other file formats by updating the file selection dialog and the corresponding data loading functions to handle different data types.

#### 2.2.2. User-Defined Analysis Settings

The user needs to enter the sampling frequency into a dialog box for the particular dataset being analyzed.

#### 2.2.3. Sliding Window Analysis

Sliding windows without overlap are generated based on the original recording, each spanning one-minute. Recordings can be as short as three minutes. However, the software has been extensively used with recordings that are at least two hours in duration, with a sampling frequency of 5000 Hz. Users can adjust the window size in the source code for shorter recordings.

#### 2.2.4. Time-Frequency Representation

During the initial processing, a spectrogram is generated for every one-minute window utilized as an input signal. 90% of the signal’s sampling frequency, used as the spectrogram window length. 80% of the signal’s sampling frequency, used as the spectrogram window overlap. The spectrogram is calculated across frequencies spanning from 1 Hz to 20 Hz, with intervals of 0.1 Hz. The calculated parameters of the spectrogram include the associated frequencies (F), the corresponding times (T), and the power spectral density estimate (P). Users can modify the spectrogram’s parameters within the source code according to their analysis. See **Figure 1-1A**.

#### 2.2.5. Identifying Theta-Dominant Segments

The relative intensity of brainwaves across various frequency ranges is assessed to measure the distribution of power in different frequency bands on the spectrogram. To compute the relative power of each band, the power spectral density values within specific frequency ranges are summed and then divided by the total sum of all power spectral density values. Specifically, the delta band’s power is calculated by summing values within the 1 to 4 Hz range, the theta band’s by summing within 5 to 15 Hz, and a higher frequency band, summing within 16 to 19 Hz. Theta dominant segments within a one-minute window are pinpointed by comparing the ratios of these relative powers, including the theta-to-delta and theta-to-higher relative power ratios, both exceeding a threshold of 1.5 and lasting for a duration of at least 5 seconds. These values can easily be changed within the code. A small caveat is that if a theta dominant segment happens to occur across the minute ‘boundary’, it could be missed – however, this is expected to be a rare occurrence and using overlapping windowing would mitigate this limitation. See **Figure 1-1B**.

#### 2.2.6. Extracted Features from Theta-Dominant Segments

To better approximate characteristics of theta power spectrum using Gaussian distribution curve, those frequencies outside the 4-13 Hz band were filtered out using bandpass filtering. The bandpass filter is created using the FIR1 function, with a specified filter order of 800. The filter’s first cutoff frequency is set at 4, while the second cutoff frequency is set to 13. A Hamming window vector of length N+1, where N represents the filter order, is employed to compute the FIR filter coefficients via the fir1 function. Following the application of the filter, the filtered signal undergoes Fast Fourier Transform (FFT) computation, enabling the derivation of its power spectral density (PSD) and the extraction of its power spectrum. The function proceeds by fitting a Gaussian curve to the power spectrum, thereby identifying peaks within the curve along with their corresponding amplitudes, frequencies, and bandwidths. A Two-Term Gaussian Model is fitted to the power spectrum.

To fit a Gaussian model to the power spectrum, we used the fit function in MATLAB. This function fits a specified model to the data, and in our case, the model was a double Gaussian (‘gauss2’). The resulting gaussian Model contained the parameters for the two Gaussian components: amplitudes (a1, a2), centers (b1, b2), and standard deviations (c1, c2). A theoretical Gaussian model was then constructed based on the fitted parameters. Finally, to identify significant peaks in the Gaussian model, we utilized the findpeaks function. This function facilitated the extraction of peak amplitudes, frequencies, bandwidths, and prominence of the identified peaks. The peak frequencies were cross-referenced with their respective power values to obtain the peak amplitudes at those frequencies.

Next, the following characteristics are extracted: The onset and termination times of the identified segment, number of peaks in power spectrum, amplitude of the peaks, frequency at which the peaks occur, and bandwidth of the peaks - width of a peak as the distance between the points where the descending signal intercepts a horizontal reference line (reference line: one-half the peak height). The extracted characteristics of each identified theta-dominant segment are then presented in a tabular format. The user can choose to download the extracted features in an Excel file format (.xlsx).

#### 2.2.7. Software Performance Evaluation using Synthetic Signal

For validation of software performance, a synthetic signal was generated over a 30-minute duration, beginning with brown noise, which was normalized to a range of -1 to 1. At the 20-minute mark, we introduced a theta oscillation at 9 Hz for 10 seconds, represented by a low-amplitude sine wave (0.1), followed by delta waves at 2 Hz for one minute, with a higher amplitude of 0.2.

In **Figure 2**, we show a synthetic signal created to test the software’s capabilities, and in **Table 1** we present the characteristics of the theta-dominant segment identified by the software. This segment matches the synthetic data, which displays theta waves for 10 seconds, starting at minute 20, with a frequency of 9 Hz. The characteristics listed in the Table confirm that our methodology accurately captures key features of the synthetic data, including the start and end times, the number of peaks, and their frequencies. This validation demonstrates that our technique effectively isolates and analyzes specific frequency components within the signal.

**Table 1:** Characteristics of the Identified Theta-Dominant Episode in Synthetic Data.

| Start Time (min) | End Time (min) | Number Of Peaks In Power Spectrum | First Peak Amp In Power Spectrum | Frequency Of First Peak In Power Spectrum Hz | Bandwidth Of First Peak In Power Spectrum Hz | Second Peak Amp In Power Spectrum | Frequency Of Second Peak In Power Spectrum Hz | Bandwidth Of Second Peak In Power Spectrum Hz |
| --- | --- | --- | --- | --- | --- | --- | --- | --- |
| 20.00 | 20.16 | 1 | 0.0025 | 9.00 | 0.33 | - | - | - |

**Figure 2:**
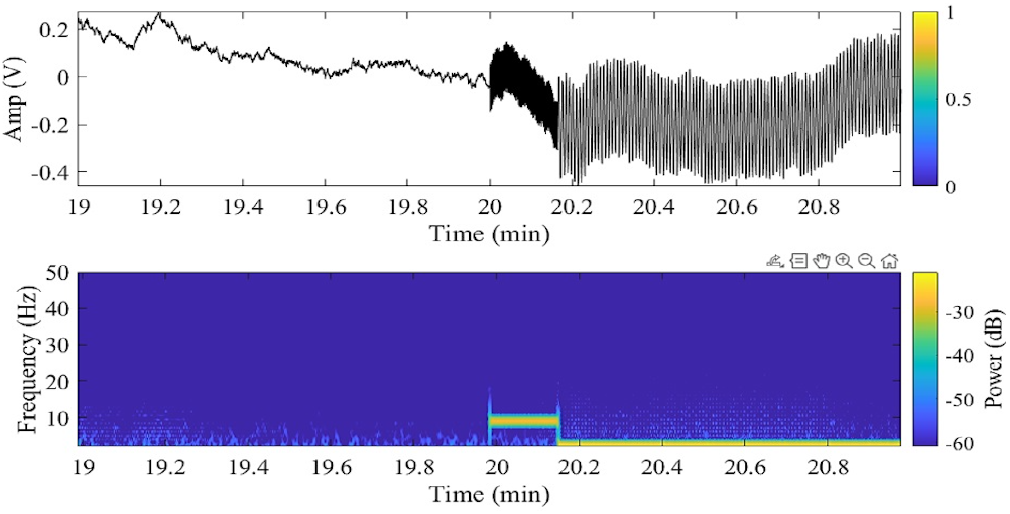
Synthetic theta waves at 9 Hz over 10 seconds, starting at minute 20. The top panel displays the signal in the time domain, illustrating its amplitude variations over time. The bottom panel presents the time-frequency representation of the signal, revealing its frequency content and how it evolves across the time axis.

### 2.3. Code Accessibility

The code and software mentioned in the paper, along with example datasets (ABF and EDF types) are available via a public OSF site [https://osf.io/rf3mn/].

## 3. Results

We illustrate our developed software using experimental data from baseline and spontaneous recurrent seizures (SRS) in a mouse model of extended hippocampal kindling (Liu et al., 2021). In **Figure 3**, we show a GUI output of the software. The example is from an SRS state for mouse C#7. A corresponding baseline state is shown in **Figure 3-1**. In **Figure 3**, the GUI output shows the raw signal of an uploaded LFP/EEG recording **(A)**, identified dominant theta activity regions in the whole recording **(B)**, as well as visual representations of spectrograms **(C)**, ratios **(D)**, identified segments **(E)**, and power spectra **(F)** corresponding to one of the identified theta-dominant segments. A Table **(G)** of all of the identified theta-dominant segments from the uploaded signal can be downloaded. To show the full extent of the features from the extracted theta-dominant segments, the downloaded table for the example of **Figure 3** is provided in **Table 2**.

**Table 2:** Event Table from Figure 3. Event table (from *Figure 3G*) is shown in full so that all columns are visible. Note that for visual purposes, the number of digits has been rounded.

| Start Time (min) | End Time (min) | Number Of Peaks In Power Spectrum | First Peak Amp In Power Spectrum | Frequency Of First Peak Amp In Power Spectrum Hz | Bandwidth Of First Peak In Power Spectrum Hz | Second Peak Amp In Power Spectrum | Frequency Of Second Peak Amp In Power Spectrum Hz | Bandwidth Of Second Peak In Power Spectrum Hz |
| --- | --- | --- | --- | --- | --- | --- | --- | --- |
| 54.91 | 55 | 2 | 0.0124 | 9 | 1.213 | 0.0066 | 12.36 | 4.67 |
| 55.01 | 55.12 | 2 | 0.0227 | 9.84 | 1.532 | 0.0098 | 12.74 | 1.204 |
| 55.16 | 55.29 | 2 | 0.0115 | 8.93 | 0.402 | 0.0052 | 9.92 | 3.135 |
| 108.73 | 108.82 | 2 | 0.0055 | 6.56 | 1.13 | 0.0045 | 8.7 | 1.169 |
| 109.66 | 109.75 | 1 | 0.0102 | 6.56 | 1.353 |  |  |  |

**Figure 3:**
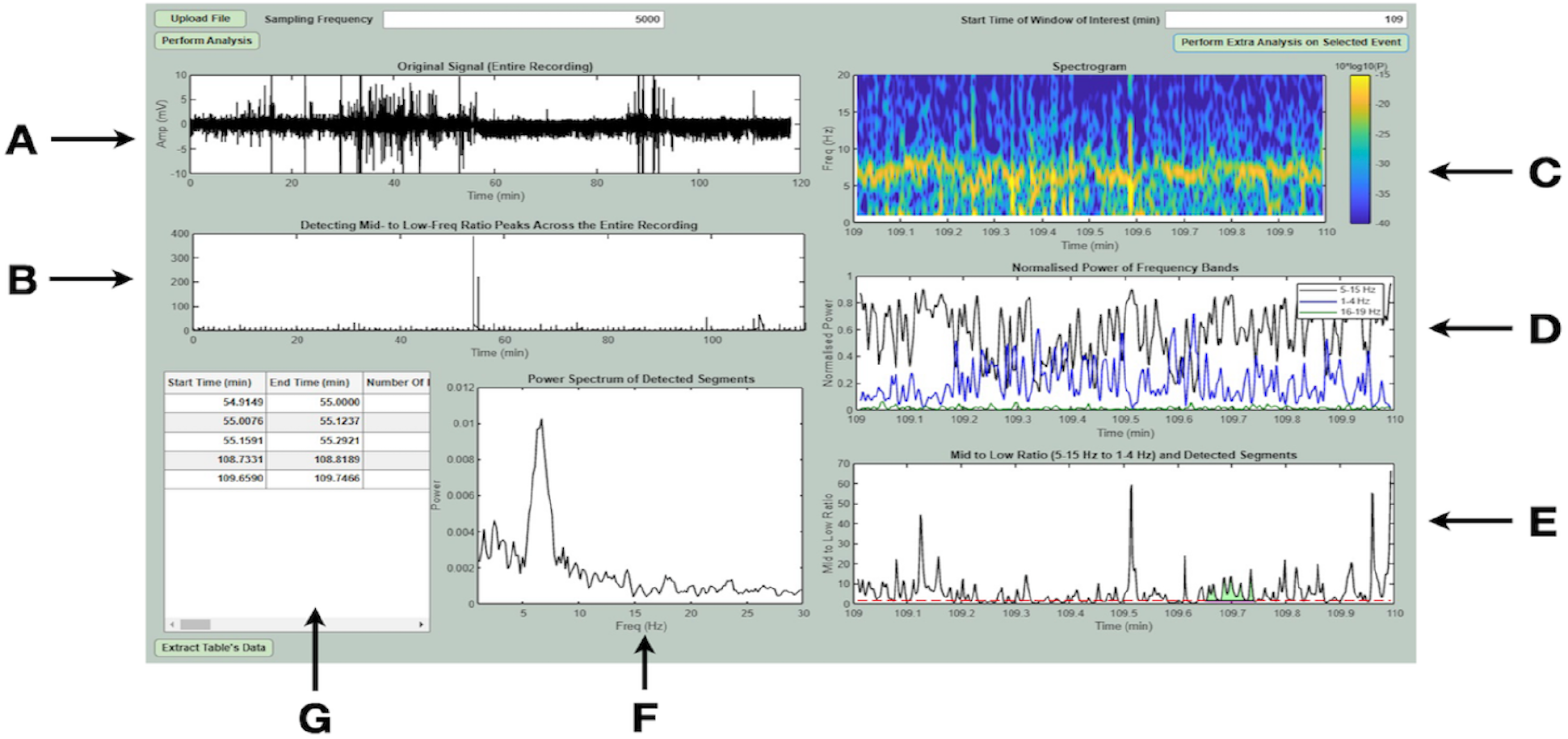
GUI Output Illustration. A comprehensive analysis of an EEG signal, focusing on the detection of theta-dominant segments. The dataset used for this example is from mouse C#7 after it has been kindled. The key components are: **(A)** *Original Signal*: Displays the raw EEG signal amplitude over time that the user has uploaded. **(B)** *Ratio Peaks*: Indicates the peak ratio (of mid-to low-frequency band) versus the starting minute of each window across the entire recording. These peak ratios include all peaks without considering whether they meet a specified ratio and duration for theta dominance or not. **(C)** *Spectrogram*: Shows the frequency content of the signal over a selected time window, highlighting changes in frequency bands. **(D)** *Normalized Powers*: Compares the normalized powers of a low-band of 1-4 Hz that includes *delta* (blue), a mid-band of 5-15 Hz that includes *theta* (black), and a high-band of 16-19 Hz (green), within the selected time window. **(E)** *Detected Segments:* Plots the theta-to-delta power ratio over time, marking the detected segments with a high enough theta activity (ratio greater than 1.5). It is shaded in green if it exceeds the specified duration (5 sec as used here) to be considered a theta dominant segment. **(F)** *Power Spectrum of Detected Segment:* Plots the power spectrum of the first detected theta-dominant segment within the selected one-minute window (indicated by the green shading). **(G)** *Event Table:* Lists the start and end times of each detected theta-dominant segment along with their corresponding characteristics. See *Table 2* for the entire table contents for this uploaded recording. **C, D**, and **E** refer to the selected time window of one-minute duration starting at the time entered (109 is specified in this example). A comparison of GUI outputs from a mouse in two different states in shown in *Figure 3-1*.

In the example of **Figure 3**, the selected one-minute window from the uploaded recording starting from minute 109 has its spectrogram displayed in **Figure 3** and the corresponding normalized powers in **Figure 3D**. In this selected minute, there is only one identified theta-dominant segment (green shading) that meets the criteria of having a minimum duration of 5 seconds and theta-delta and theta-higher ratios exceeding 1.5. The power spectrum of this segment is shown in **Figure 3F** and can be seen to have a single peak.

In **Figure 4**, we show a power spectrum with a double peak, corresponding to an identified theta-dominant segment from an earlier part of the same uploaded dataset of **Figure 3**. In **Table 2**, we can see the particular frequencies, powers and bandwidths of this theta-dominant segment with two peaks, as well as all the other identified theta-dominant segments in this two-hour recording.

**Figure 4:**
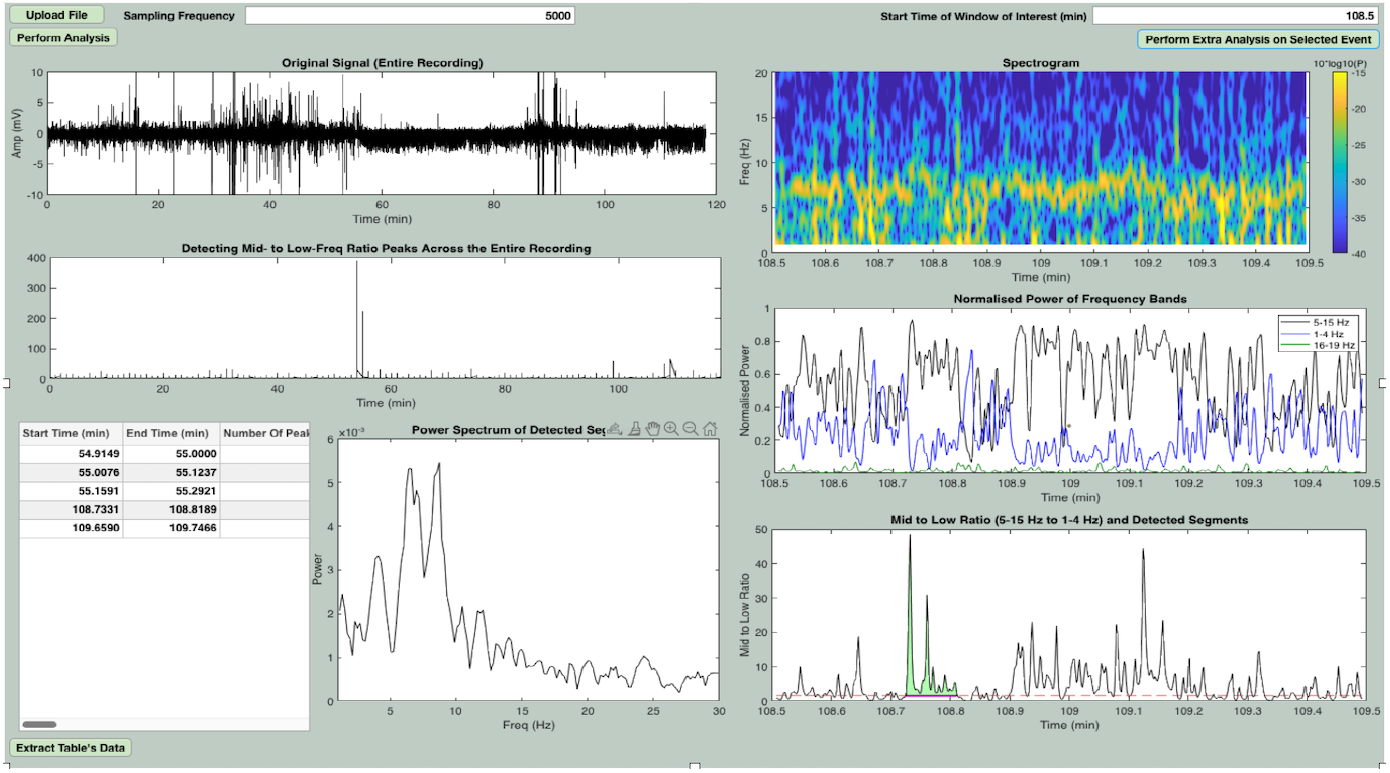
Double Peak Power Spectra in Selected Theta-Dominant Segment. The same dataset from *Figure 3* is used but a different one-minute time window (starting at 108.5) is used to show a theta-dominant segment that has a double peak.

To better appreciate how the threshold ratios ‘translate’ to power spectra, in **Figure 5**, we expand on the identified theta-dominant segment shown in **Figure 3**. If we compare the theta, delta and higher frequency peaks of the power spectra of this identified segment, we see that the theta peak is about twice the size of the delta one and is clearly dominant.

**Figure 5:**
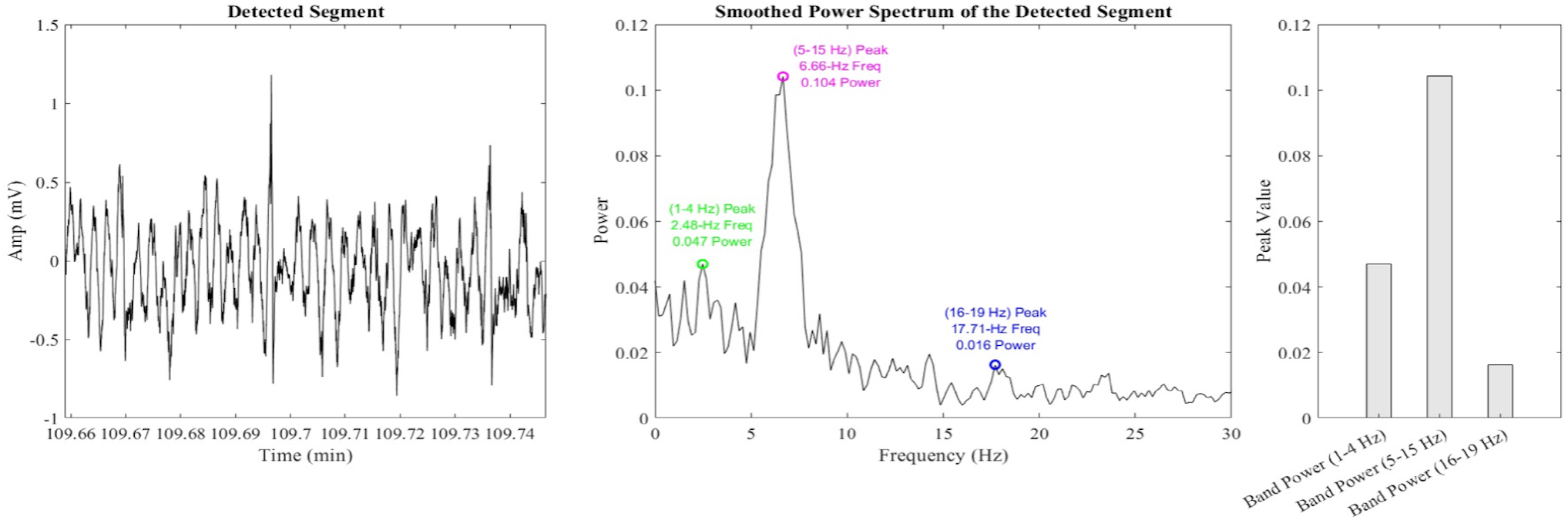
Analysis of a Theta-Dominant Segment. The left panel shows the original theta-dominant segment waveform over a 5-second interval (part of an identified segment that has a duration of 5.4 s). The dataset used is the same as in *Figure 2*. The middle panel displays the smoothed power spectrum of the 5.4 s duration theta-dominant segment, showing the peaks at low (delta) (1.48 Hz, 0.055 power), mid (theta) (7.59 Hz, 0.087 power), and high (18.33 Hz, 0.017 power) frequencies. The right panel compares the peak power values for the low-, mid- and high-frequency bands, highlighting the dominance of the theta (mid-) peak.

In **Figure 6** we show an expansion of the identified segment along with its corresponding spectrogram in the selected minute. To illustrate how one’s choice of minimum duration and ratio affects the identification of theta-dominant segments, we show in **Figure 7** how the identified segments (green shading) in this particular minute would change if these parameter values are adjusted. Clearly, the stricter one is with the duration and ratio, the less likely that theta-dominant segments would be identified. That is, none would have been found in this selected minute if the minimum duration was 6 seconds and the ratio threshold was 3. On the other hand, if a minimum duration of 2 seconds was used with a threshold of 1, then many theta-dominant segments would have been identified.

**Figure 6:**
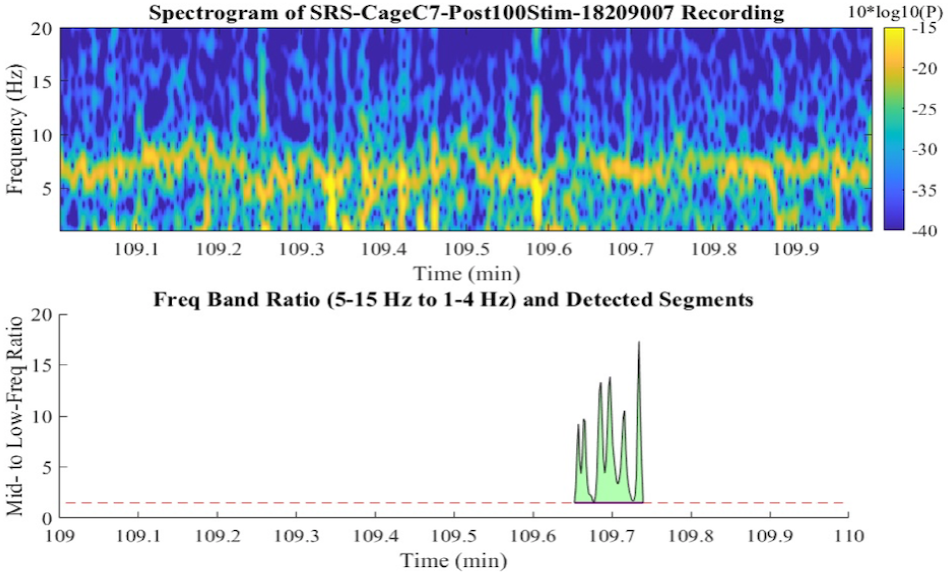
Spectrogram and Theta-Delta Ratio Analysis. The top panel displays a spectrogram of a signal over a one-minute period, with frequency (Hz) on the y-axis and time (min) on the x-axis. The color intensity represents the power spectral density, with warmer colors indicating higher power. The bottom panel shows the theta-delta ratio over the same time period, with a detected segment highlighted in green. The ratio peaks around 109.7 min, indicating a significant increase in theta power relative to delta power during this interval. The dataset used is the same as in *Figure 3*.

**Figure 7:**
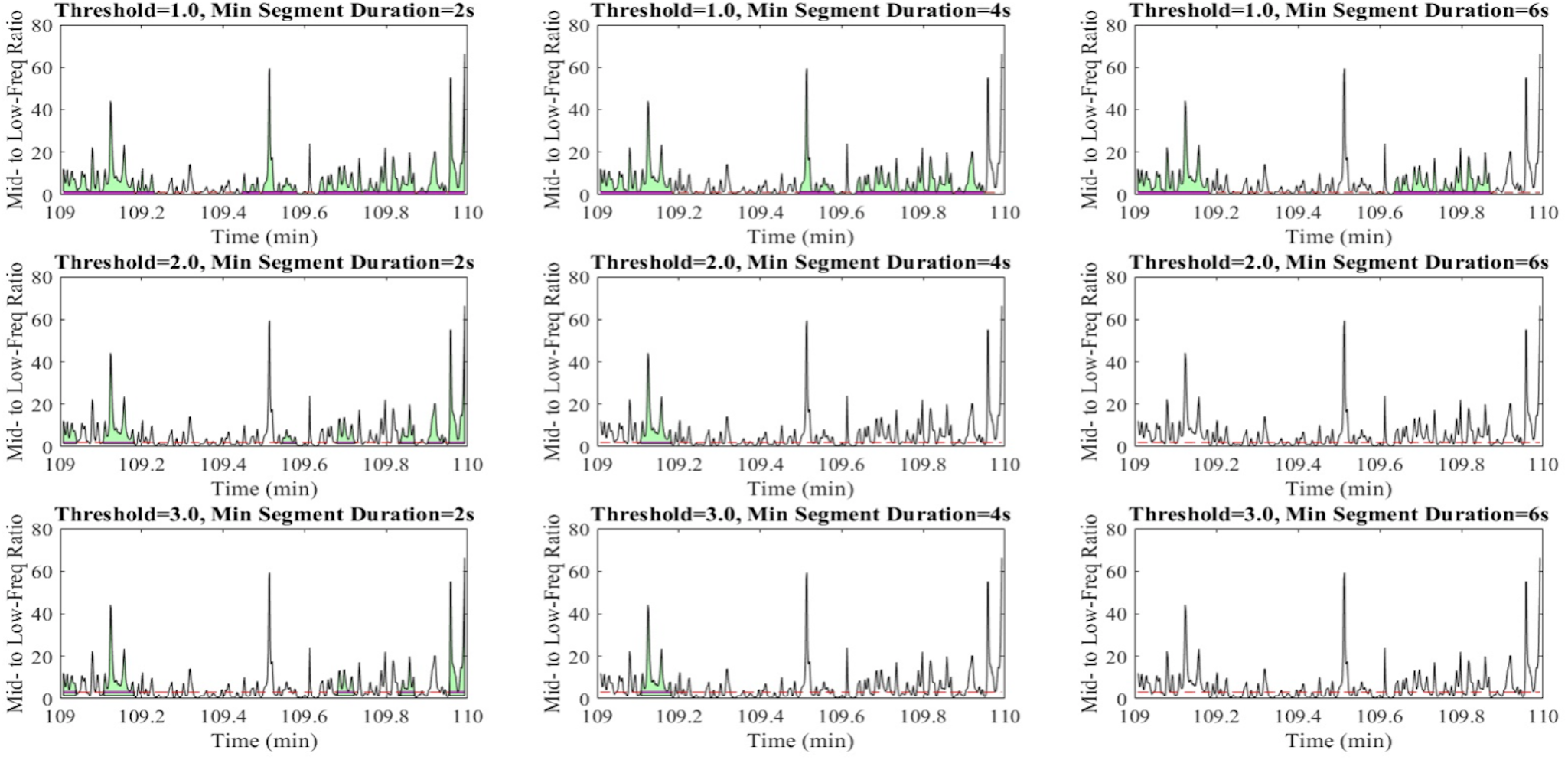
Theta Dominance Detection Illustrations Using Different Thresholds and Minimum Durations. The figure shows nine subplots depicting the theta-delta ratio over a one-minute period where the thresholds and minimum duration choices are changed to values as indicated. The rows represent different thresholds (1.0, 2.0, and 3.0), and the columns represent different minimum durations (2s, 4s, and 6s). Green shaded regions indicate detected theta-dominant segments where the theta-delta ratio exceeds the threshold for at least the specified duration. This is the same dataset and range as shown in *Figure 6* where a threshold of 1.5 and a minimum duration of 5s was used. As both threshold and minimum duration increase, fewer segments are detected, indicating stricter criteria for theta-dominant segment identification, so that at the largest threshold and duration shown, no theta-dominant segments are detected.

To illustrate a detailed analysis of data using our software, we analyzed many hours of baseline and SRS state data for two mice. In general, since the baseline theta-dominant segments were much smaller than the SRS ones (see **Figure 3-1**), we removed those that had amplitudes less than 1e-4. In **Table 3**, a comparison between baseline (B) and SRS states in terms of the number of theta-dominant segments and the means and variances of their characteristics can be seen. Note that the percent of time for which theta dominance occurs in the many hours of recordings is small (4^th^ column), thus reflecting the challenge of manually finding robust theta activity in long recordings. Further, because of being able to analyze many hours of data, statistical comparisons of the extracted features of theta activities can be performed. Specifically, we find that there are statistically significant differences in the frequencies and amplitudes between baseline and SRS states (Mann-Whitney nonparametric test, considered significant if p<0.05; Freq: p=0.005 for C#7, p=0.004 for F#17; Amp: p=1.02e-20 for C#7, p=1.07e-11 for F#17). There are not statistically significant differences in the durations (p=0.342 for C#7, p=0.068 for F#17), and only for C#7 is there a significant difference in the bandwidth (p=6.36e-8 for C#7, p=0.22 for F#17).

**Table 3:**
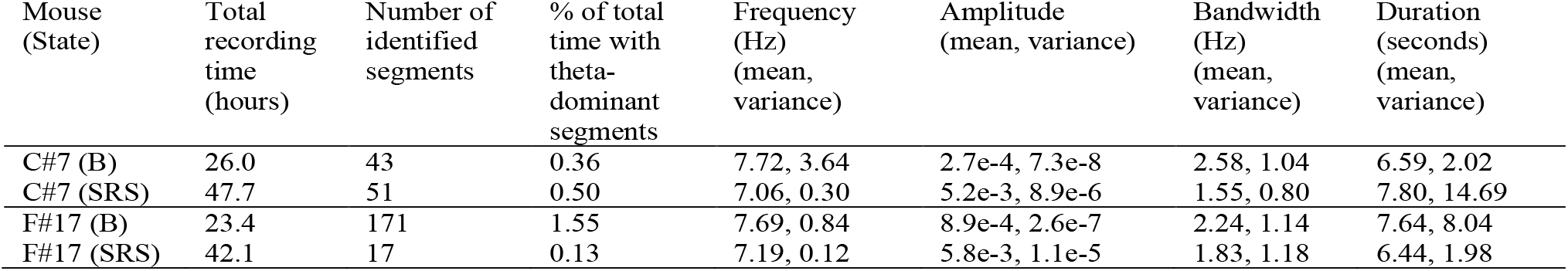
Comparison Between Baseline (B) and SRS States in Two Mice. The values for frequencies, amplitudes and bandwidths refer to the lower theta frequency (peak 1) if there were more than one peaks obtained from the fit.

| Mouse (State) | Total recording time (hours) | Number of identified segments | % of total time with theta-dominant segments | Frequency (Hz) (mean, variance) | Amplitude (mean, variance) | Bandwidth (Hz) (mean, variance) | Duration (seconds) (mean, variance) |
| --- | --- | --- | --- | --- | --- | --- | --- |
| C#7 (B) | 26.0 | 43 | 0.36 | 7.72, 3.64 | $2.7e-4$, $7.3e-8$ | 2.58, 1.04 | 6.59, 2.02 |
| C#7 (SRS) | 47.7 | 51 | 0.50 | 7.06, 0.30 | $5.2e-3$, $8.9e-6$ | 1.55, 0.80 | 7.80, 14.69 |
| F#17 (B) | 23.4 | 171 | 1.55 | 7.69, 0.84 | $8.9e-4$, $2.6e-7$ | 2.24, 1.14 | 7.64, 8.04 |
| F#17 (SRS) | 42.1 | 17 | 0.13 | 7.19, 0.12 | $5.8e-3$, $1.1e-5$ | 1.83, 1.18 | 6.44, 1.98 |

From our many hours of analyses, we can also consider when and where theta-dominant segments occur relative to ictal events in the SRS states. For example, in **Figure 8**, we show plots for the two mice of their frequencies over continuous time. It can be seen that dominant theta activities can occur just before an ictal event, but theta-dominant segments are not detected for many minutes post-ictal after the seizure – for these two mice, it was at least 20 minutes before a theta-dominant segment could be detected after a seizure occurrence.

**Figure 8:**
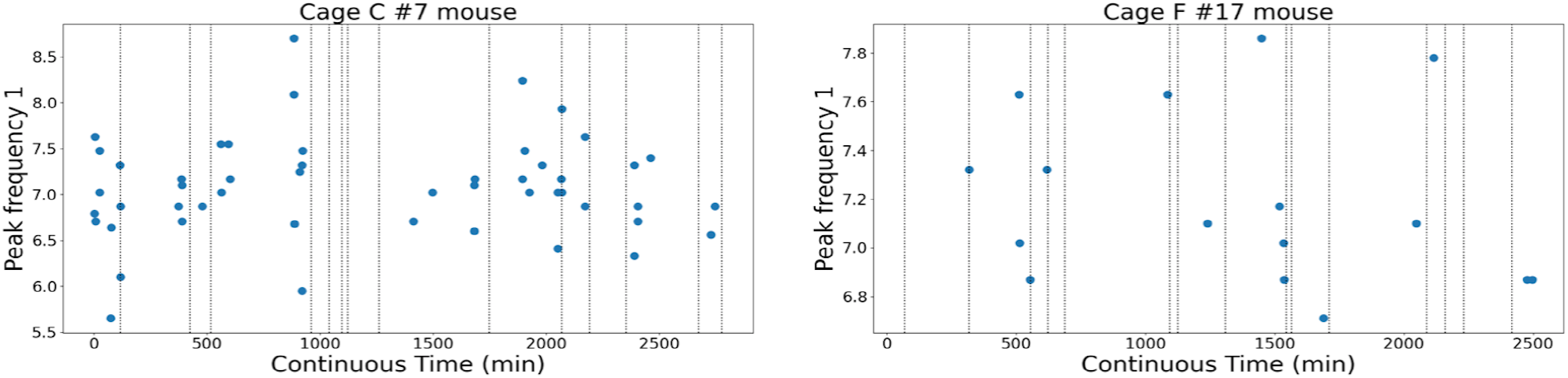
Theta-dominant Segment Timing Relative to Ictal Event Discharges. The dashed lines represent the time of ictal discharge occurrences. Blue filled circles indicate identified theta-dominant segments. The (first) peak frequency (Hz) feature for mouse C#7 is shown on the left and for mouse F#17 on the right. The x-axis represents continuous time during the SRS state. Note that if the blue dot looks super-imposed on the dashed line (ictal discharge occurrence), it is very close to the start of the ictal discharge and cannot be discerned in the plot resolution. Also, sometimes there are clusters of seizures in the datasets.

To ensure that our exact filter choices did not play a critical role in our results, we examined the impact of filtering by analyzing theta-dominant episodes from one of the SRS recordings. We compared the characteristics of 14 theta-dominant segments with and without applying zero-phase digital filtering and using different filter orders. This is shown in **Figure 9** for the frequency of the first peak.

**Figure 9:**
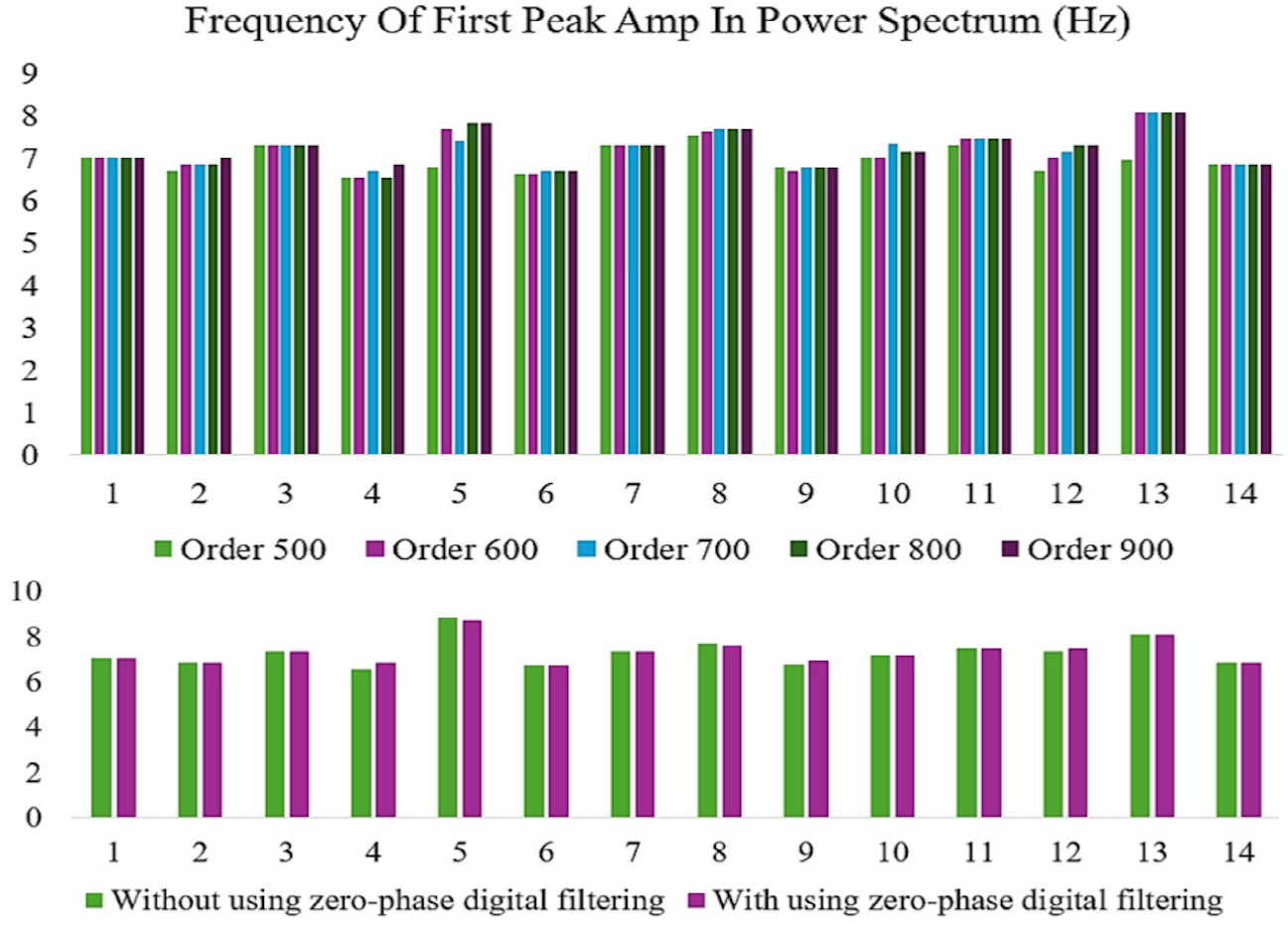
Comparison of theta-dominant episodes with and without zero-phase filtering and different filter orders. The figure illustrates this for the frequency of the first peak amplitude. Different filter orders are shown at the top and with and without zero-phase filtering is shown on the bottom. Analysis of first peak amplitude, frequency, and bandwidth of the power spectrum revealed no significant differences between the two conditions (p-values: amplitude = 0.9958, frequency = 0.9018, bandwidth = 0.4378), indicating that zero-phase filtering does not significantly impact these characteristics. Analysis of the first peak amplitude frequency in the power spectrum for various filter orders (500 to 900) showed no statistically significant differences, as confirmed by t-tests. Specifically, the p-values for comparisons between adjacent filter orders— Order 500 vs. Order 600 (p = 0.206), Order 600 vs. Order 700 (p = 0.798), Order 700 vs. Order 800 (p = 0.911), and Order 800 vs. Order 900 (p = 0.843)—were all above the conventional threshold of 0.05.

**Figure 10:**
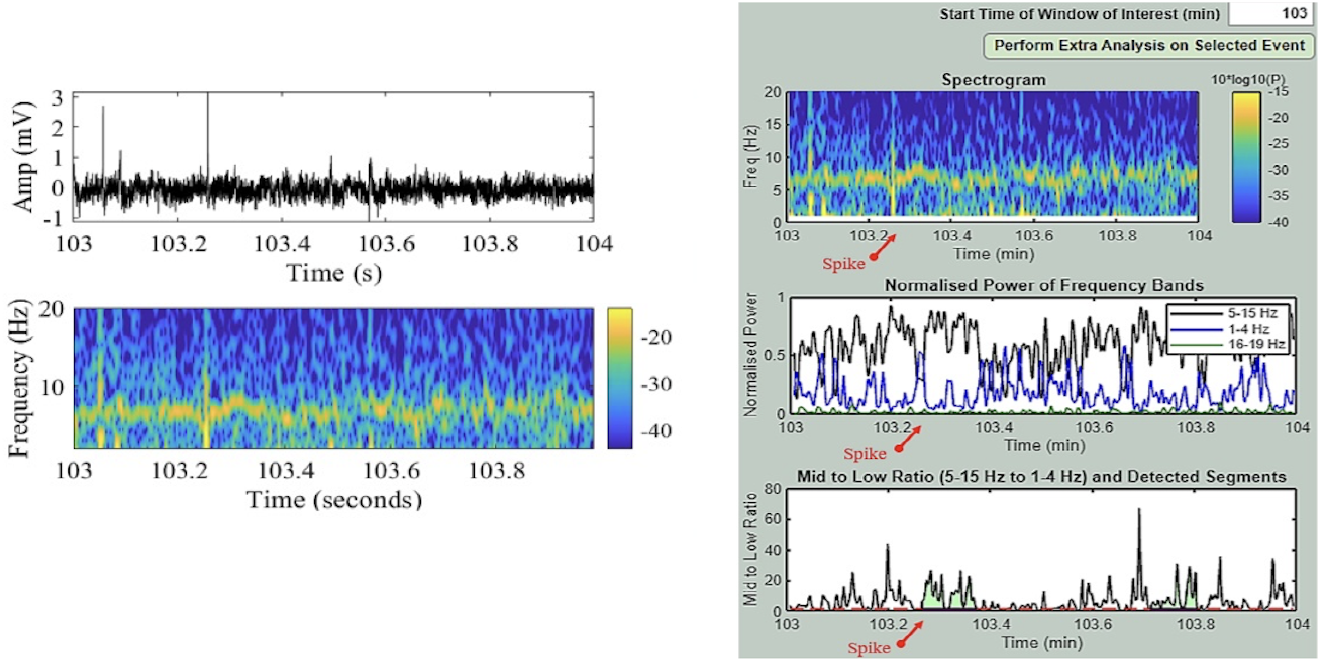
Disruption of Theta Continuity by Spikes. Recording number 18206001 from Cage C #7, minute 103, SRS state (post 100 stimulations). (Left) The original signal along with its corresponding spectrogram. (Right) The output from the software. When interictal spikes occur, the segments of theta activity that precede and follow these spikes can be considered distinct episodes, provided they meet the specified criteria for a theta-dominant segment.

Spikes can disrupt the continuity of theta activity (see **Figure 10** (left)), leading to the segmentation of theta episodes. When spikes occur, the segments of theta activity before and after these spikes are treated as separate episodes if they meet the specified criteria for a theta-dominant segment. When we refer to “theta dominant”, it implies that there is a continuous presence of theta activity without interruptions. Continuous theta indicates a sustained state of theta rhythm. Interruptions from spikes or other activities can lead to disruptions in that continuity.

## 4. Discussion

Population oscillations are omnipresent and are fundamental to our understanding of brain mechanisms (Buzsáki, 2006). Theta frequency rhythms in the hippocampus are one of the most heavily studied rhythms due to their association and modulation with learning, memory and pathologies. However, a combination of experimental variability and non-standardized experimental analyses make it difficult to understand the expression and modifications of theta rhythms in different states. To overcome this, we have developed an easy-to-use advanced software tool that can extract dominant theta activities and characterize their features. In this paper, we presented our software and illustrated its use with normal and pathological datasets of seizures in epilepsy. We showed that statistically significant differences in theta features between baseline and seizure states can be obtained, and that the appearance of theta activities after a seizure can be quantified. Specifically, there was a significant difference in both frequency and power for two different mice, and a much larger time post-ictal compared to pre-ictal was required for dominant theta activities. We will present a full and extensive analysis of more datasets in a future publication.

Given the range and variability of experimental data associated with theta (and other) rhythms, it is of utmost importance to be able to analyze data in a standardized manner so that comparisons can be robustly done. Rather than manually selecting time windows (Tort et al., 2018; Lopes-dos-Santos et al., 2018; Kienitz et al., 2021; Tort, Brankačk, & Draguhn, 2018, Song et al., 2024) in which theta rhythms occur, the user can automatically extract dominant theta activities in a standardized fashion once a relative ratio (to delta and higher frequency rhythms) threshold and minimal duration is set. Not only would this streamline analyses and translate into huge time savings, but also, it would allow the user to monitor and track changes in rhythms in a quantified, feature-specific fashion. In turn, this would enable these rhythms to potentially serve as disease biomarkers and evaluators of drug testing and usage.

The theta rhythms showed a frequency band with temporal variability, occasionally displaying dual peaks in frequency distribution. This variability illustrates that the notion of a single-peak phenomenon for theta rhythms should not be assumed during analysis, as might be expected given that two types of theta activities have long been recognized (Kramis et al., 1975). For example, we found that the theta-dominant segments obtained for the two mice (**Table 3**) occasionally displayed dual peaks in frequency distribution (approximately one-third of the time), and was different for baseline and SRS states, suggesting that this could be a feature of focus for seizure states. With enough data, a full analysis may reveal that there are consistent differences in frequency distributions across consecutive theta-dominant segments, between pre-ictal states, post-ictal states, seizure severity, inter-ictal spike presence and seizure-cluster dependence. In general, a monitoring of changes in theta rhythm features can be undertaken.

Being able to quantify rhythmic changes in a robust and feature-specific fashion will be critically helpful for mathematical modelers of brain rhythms. Theoretical and mathematical network models are essential to help us determine underlying brain mechanisms that give rise to population rhythms. While there has been extensive modeling of these rhythms (Ferguson and Skinner, 2022), the best choices and appropriateness of different models being developed is not obvious and usually not clear (Levenstein et al., 2023). Knowing what oscillation features change robustly during disease could provide another level of constraint in designing and using mathematical model to help uncover brain mechanisms.

In conclusion, a novel and easy-to-use software that effectively identifies dominant theta rhythms has been developed. This dominance is defined in relation to delta waves for a minimal duration in time. Experimental data confirmed the software’s efficacy and significant theta feature differences between normal and pathological states have been shown to exist after analyzing many hours of recordings.

## Extended Figures

**Figure 1-1:**
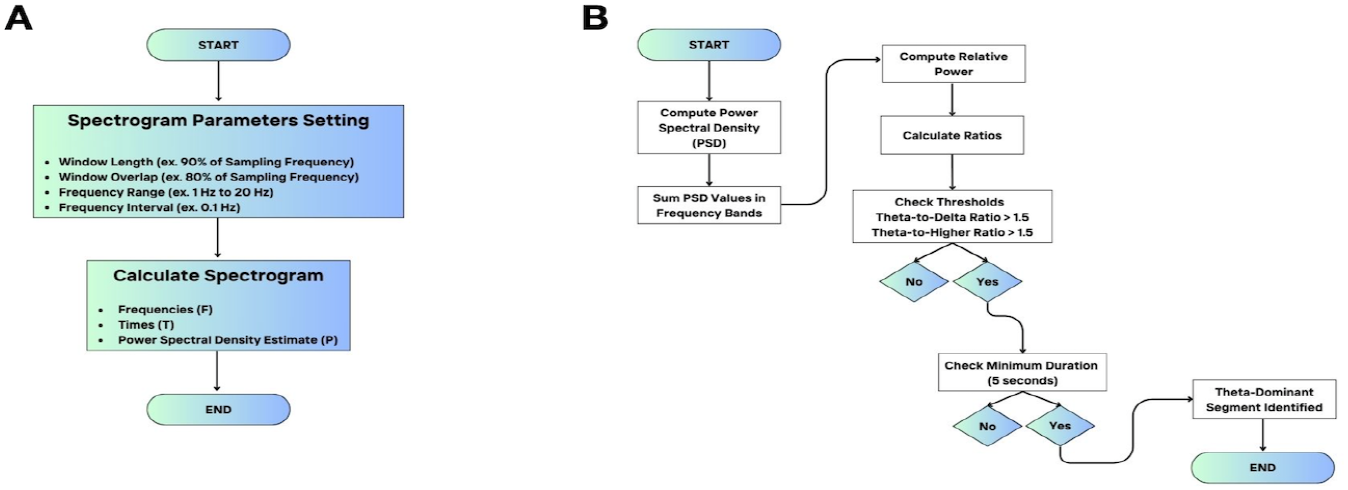
Overviews of parts of the Neuro Dominance Tracker Software. An overview of: **(A)** the generation of spectrograms (as described in section 2.2.4 of the Methods), and **(B)** the identification of theta-dominant segments (as described in section 2.2.5 of the Methods).

**Figure 3-1:**
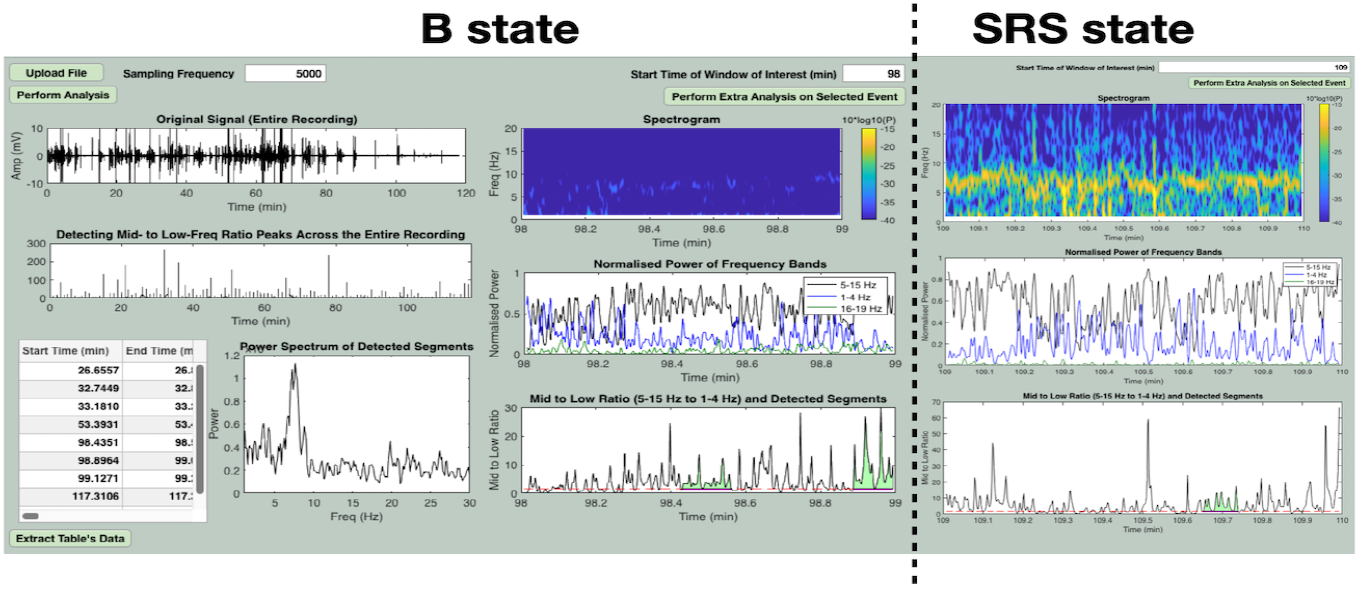
GUI Outputs from a Mouse in Different States. A dataset from a mouse (C#7) in a baseline (B) state (left side) and in a kindled state in which there are spontaneous recurrent seizures (SRS). The same dataset as in *Figure 3G* is used for the SRS state in illustrating these output differences, but only part of the GUI is shown. From these outputs, it is clear that the baseline theta-dominant segments are lower amplitude (compare spectrograms that have same range) and that there are more of them (compare Event Table here for B state, and that in *Figure 3G* for SRS state). Also note that the baseline state shows two theta-dominant segments for the selected one-minute window.

## References

Bin, N.-R., Song, H., Wu, C., Lau, M., Sugita, S., Eubanks, J. H., & Zhang, L. (2017). Continuous Monitoring via Tethered Electroencephalography of Spontaneous Recurrent Seizures in Mice. Frontiers in Behavioral Neuroscience, 11, 172. 10.3389/fnbeh.2017.00172

Buzsáki, G. (2002). Theta oscillations in the hippocampus. Neuron, 33(3), 325–340.

Buzsaki, G. (2006). Rhythms of the Brain (1st ed.). Oxford University Press, USA.

Chauvière, L., Rafrafi, N., Thinus-Blanc, C., Bartolomei, F., Esclapez, M., & Bernard, C. (2009). Early deficits in spatial memory and theta rhythm in experimental temporal lobe epilepsy. The Journal of Neuroscience: The Official Journal of the Society for Neuroscience, 29(17), 5402–5410. 10.1523/JNEUROSCI.4699-08.2009

Colgin, L. L. (2013). Mechanisms and Functions of Theta Rhythms. Annual Review of Neuroscience, 36(1), 295–312. 10.1146/annurev-neuro-062012-170330

Colgin, L. L. (2016). Rhythms of the hippocampal network. Nature Reviews Neuroscience, 17(4), 239–249. 10.1038/nrn.2016.21

Fedor, M., Berman, R. F., Muizelaar, J. P., & Lyeth, B. G. (2010). Hippocampal Theta Dysfunction after Lateral Fluid Percussion Injury. Journal of Neurotrauma, 27(9), 1605–1615. 10.1089/neu.2010.1370

Ferguson, K.A., Skinner, F.K. (2022). Hippocampal Theta, Gamma, and Theta/Gamma Network Models. In: Jaeger, D., Jung, R. (eds) Encyclopedia of Computational Neuroscience. Springer, New York, NY. 10.1007/978-1-0716-1006-0_27

Franklin KBJ, Paxinos G. (1997). The mouse brain in stereotaxic coordinates. San Diego, CA: Academic Press.

Goutagny, R., Gu, N., Cavanagh, C., Jackson, J., Chabot, J.-G., Quirion, R., Krantic, S., & Williams, S. (2013). Alterations in hippocampal network oscillations and theta–gamma coupling arise before Aβ overproduction in a mouse model of Alzheimer’s disease. European Journal of Neuroscience, 37(12), 1896–1902. 10.1111/ejn.12233

Kienitz R, Cox MA, Dougherty K, Saunders RC, Schmiedt JT, Leopold DA, et al. (2021). Theta, but Not Gamma Oscillations in Area V4 Depend on Input from Primary Visual Cortex [Internet]. Vol. 31, Current Biology. Elsevier BV; p. 635–42.e3. Available from: 10.1016/j.cub.2020.10.091.

Kramis, R., Vanderwolf, C. H., & Bland, B. H. (1975). Two types of hippocampal rhythmical slow activity in both the rabbit and the rat: Relations to behavior and effects of atropine, diethyl ether, urethane, and pentobarbital. Experimental Neurology, 49(1), 58–85. 10.1016/0014-4886(75)90195-8

Levenstein, D., Alvarez, V. A., Amarasingham, A., Azab, H., Chen, Z. S., Gerkin, R. C., Hasenstaub, A., Iyer, R., Jolivet, R. B., Marzen, S., Monaco, J. D., Prinz, A. A., Quraishi, S., Santamaria, F., Shivkumar, S., Singh, M. F., Traub, R., Nadim, F., Rotstein, H. G., & Redish, A. D. (2023). On the Role of Theory and Modeling in Neuroscience. Journal of Neuroscience, 43(7), 1074–1088. 10.1523/JNEUROSCI.1179-22.2022

Lisman, J., Buzsáki, G., Eichenbaum, H., Nadel, L., Rangananth, C., & Redish, A. D. (2017). Viewpoints: How the hippocampus contributes to memory, navigation and cognition. Nature Neuroscience, 20(11), 1434–1447. 10.1038/nn.4661

Liu H, Tufa U, Zahra A, Chow J, Sivanenthiran N, Cheng C, Liu Y, Cheung P, Lim S, Jin Y, Mao M, Sun Y, Wu C, Wennberg R, Bardakjian B, Carlen PL, Eubanks JH, Song H, Zhang L. (2021). Electrographic Features of Spontaneous Recurrent Seizures in a Mouse Model of Extended Hippocampal Kindling. Cereb Cortex Commun. 2021 Jan 22;2(1):tgab004. doi: 10.1093/texcom/tgab004. PMID: 34296153; PMCID: PMC8152854.

Lopes-dos-Santos V, van de Ven GM, Morley A, Trouche S, Campo-Urriza N, Dupret D. (2018). Parsing Hippocampal Theta Oscillations by Nested Spectral Components during Spatial Exploration and Memory-Guided Behavior [Internet]. Vol. 100, Neuron. Elsevier BV; p. 940–52.e7. Available from: 10.1016/j.neuron.2018.09.031..

Oostenveld, R. et al., 2011. FieldTrip: Open Source Software for Advanced Analysis of MEG, EEG, and Invasive Electrophysiological Data. Computational Intelligence and Neuroscience, 2011, pp.1–9. Available at: 10.1155/2011/156869.

Milikovsky, D. Z., Weissberg, I., Kamintsky, L., Lippmann, K., Schefenbauer, O., Frigerio, F., Rizzi, M., Sheintuch, L., Zelig, D., Ofer, J., Vezzani, A., & Friedman, A. (2017). Electrocorticographic Dynamics as a Novel Biomarker in Five Models of Epileptogenesis. Journal of Neuroscience, 37(17), 4450–4461. 10.1523/JNEUROSCI.2446-16.2017

Schultheiss, N. W., Schlecht, M., Jayachandran, M., Brooks, D. R., McGlothan, J. L., Guilarte, T. R., & Allen, T. A. (2020). Awake delta and theta-rhythmic hippocampal network modes during intermittent locomotor behaviors in the rat. Behavioral Neuroscience, 134, 529–546. 10.1037/bne0000409

Song H, Mah B, Sun Y, Aloysius N, Bai Y, Zhang L. (2024). Development of spontaneous recurrent seizures accompanied with increased rates of interictal spikes and decreased hippocampal delta and theta activities following extended kindling in mice. Exp Neurol. 2024 Sep;379:114860. doi: 10.1016/j.expneurol.2024.114860. Epub 2024 Jun 12. PMID: 38876195.

Song H, Tufa U, Chow J, Sivanenthiran N, Cheng C, Lim S, Wu C, Feng J, Eubanks JH, Zhang L. (2018). Effects of Antiepileptic Drugs on Spontaneous Recurrent Seizures in a Novel Model of Extended Hippocampal Kindling in Mice. Front Pharmacol. 2018 May 18;9:451. doi: 10.3389/fphar.2018.00451. PMID: 29867462; PMCID: PMC5968120.

Tort, A.B.L., Ponsel, S., Jessberger, J. et al. (2018). Parallel detection of theta and respiration-coupled oscillations throughout the mouse brain. Sci Rep 8, 6432. 10.1038/s41598-018-24629-z.

Tort ABL, Brankačk J, Draguhn A. (2018). Respiration-Entrained Brain Rhythms Are Global but Often Overlooked [Internet]. Vol. 41, Trends in Neurosciences. Elsevier BV; p. 186–97. Available from: 10.1016/j.tins.2018.01.007..

Song H, Mah B, Sun Y, Aloysius N, Bai Y, Zhang L. Development of spontaneous recurrent seizures accompanied with increased rates of interictal spikes and decreased hippocampal delta and theta activities following extended kindling in mice. Exp Neurol. 2024 Sep;379:114860. doi: 10.1016/j.expneurol.2024.114860. Epub 2024 Jun 12. PMID: 38876195.

